# Myeloid-mediated IL-1R signaling in immuno-responsive Thy-1 negative fibroblasts is critical for pulmonary fibrosis

**DOI:** 10.1101/2021.05.11.443647

**Authors:** Daniel Abebayehu, Chiuan-Ren Yeh, Grace C. Bingham, Sarah Ewald, Thomas H. Barker

## Abstract

Idiopathic pulmonary fibrosis (IPF) is a fatal disease with poorly defined pathogenic mechanism and no cure. It is characterized by chronic inflammation, myofibroblast accumulation, and aberrant extracellular matrix (ECM) remodeling. Fibrosis progression is considered to occur due to sustained aberrant fibroblast mechanotransduction: sensing “normal” soft tissue as stiff scarred tissue leading to the overproduction of ECM that then stiffens the microenvironment, thus reinforcing a progressive, stiffness-dependent fibrotic program. How chronic inflammation leads to aberrant mechanotransduction is not well understood. Thy-1 is a regulator of mechanotransduction in fibroblasts. Thy-1 expression is lost in fibroblastic foci, the active sites of fibrosis, although the mechanism of this loss is unknown. We demonstrate that in IPF tissue, the αSMA+ fibroproliferative foci express the Type 1 IL-1 receptor (IL-1RI) and IL-1RI-deficient mice did not develop bleomycin-induced pulmonary fibrosis. Using ASC speck formation during inflammasome activation as a marker of mature IL-1β release, we identified the immune compartment as the source of active IL-1β during bleomycin-induced fibrosis. Furthermore, incubating mouse lung fibroblasts on soft (2kPa) hydrogels with IL-1β was sufficient to reduce Thy-1 surface expression and induce αvβ3 integrin activation. As expected, Thy-1 negative fibroblasts exhibited elevated αvβ3 integrin activation but surprisingly, Thy-1 negative fibroblasts also expressed higher levels of IL-1RI, potentially linking the immuno-responsive and mechanosensitivity of this fibroblast subpopulation. Leveraging the non-resolving fibrosis that occurs in Thy-1^-/-^ mice, we observed that crossing Thy-1^-/-^ mice onto the IL-1RI^-/-^ background was sufficient to reduce fibrosis. Together, these data indicate that Thy-1 negative fibroblasts are an immuno-responsive subpopulation that also display altered mechanotransduction, potentially serving as the link between the noted inflammation and aberrant mechanotransduction observed in IPF.

## Introduction

Idiopathic pulmonary fibrosis (IPF) is a fatal disease characterized by excessive extracellular matrix (ECM) deposition within the lung interstitium. IPF has no known etiology, no cure, a high clinical burden, extensively studied inflammation and tissue remodeling, yet it remains unknown what triggers the aberrant remodeling [1,2]. IPF is characterized by chronic inflammatory cell infiltration, high levels of inflammatory markers, aberrant extracellular matrix (ECM) deposition, and the accumulation of activated fibroblasts termed ‘myofibroblasts’. As IPF progresses, the lung interstitium becomes an order of magnitude stiffer (from approximately 2kPa to 20-25kPa) [3]. However, the sites of active fibrosis are the fibroblastic foci, which consist of myofibroblasts that drive fibrosis progression [4]. Fibroblastic foci are considered the leading edge of fibrosis, directing the fibrotic destruction of the lung [5]. Previous work has characterized the fibroblastic foci has being rich in Thy-1 deficient fibroblasts [6] while possessing a tissue stiffness not dissimilar from normal lung tissue (∼2kPa) [7]. Thy-1/CD90, a glycosylphosphatidylinositol (GPI)-anchored surface protein with an integrin-binding domain, participates in cell-cell and cell-matrix interactions among T-cells, endothelial cells, neurons, mesenchymal stem cells, and fibroblasts [8,9]. Within fibroblasts, Thy-1 has been shown to contribute to mechanotransduction via cis-interaction with integrin αvβ3 and altering integrin αvβ3 conformation in a rigidity-dependent manner [10,11]. The loss of Thy-1 confers stiffness agnostic behavior to fibroblasts which promotes a myofibroblastic phenotype even while on soft (∼2kPa) substrates [7]. The environmental signals that promote the loss of Thy-1 in lung fibroblasts are currently unknown.

In addition to the tissue remodeling that occurs, IPF is also characterized by chronically elevated immune cell infiltrate and inflammatory markers [12]. There is a large body of work implicating IL-1R signaling in pulmonary fibrosis, from polymorphisms in the signaling cascade among IPF patients[13] to elevated levels of IL-1β in the fibrotic lungs of mice and IPF patients [14,15]. Intratracheal IL-1β delivery can induce pulmonary fibrosis in mice, however the mechanism of this biology is unclear [16]. While inflammation is a noteworthy component of IPF, clinical trials using anti-inflammatory therapies targeting IFNγ (simtuzumab) [17], TNFα (prednisone) [18], and IL-13 (tralokinumab and lebrikizumab) [19,20] signaling pathways have been widely unsuccessful [21], reflecting a gap in our understanding of inflammation and altered mechanotransduction in fibrotic tissue remodeling. A model consistent with these findings would be a causal role for inflammation in the onset of the disease (rather than during late stage IPF when patients are typically diagnosed and treated) followed by a stiffness-dependent hyper activation of ECM-depositing fibroblasts would explain the conflicting experimental and clinical data. Based on this premise, we hypothesize that Thy-1 loss in lung fibroblasts occurs in response to inflammatory cytokines. In this way, the Thy-1 negative subpopulation would be the bridge that responds to inflammatory cues and subsequently promoting the canonical stiffness-dependent fibrotic program as previously shown [7].

In this study, we demonstrate that within the fibrotic lesions of tissue from IPF patients, cells express both αSMA and IL-1RI, the type I IL-1 receptor for IL-1β. Using the bleomycin-induced fibrosis model, mice lacking IL-1RI demonstrated reduced fibrosis. The proteolytic processing and release of mature IL-1β depends on formation of in inflammasome signaling complexes containing ASC specks. Using ASC Spec formation as a surrogate for the initiation of IL-1β -IL-1R1 signaling, we demonstrate that immune cells within the lung are the source of IL-1β *in vivo* during bleomycin-induced fibrosis. When treated with IL-1β, mouse lung fibroblasts seeded on 2kPa hydrogels lose Thy-1 expression and display αvβ3 in an active conformation. Furthermore, Thy-1 negative fibroblasts express higher levels of IL-1RI, indicating they are uniquely equipped to be immuno-responsive. Finally, we leveraged the observation that bleomycin-induced fibrosis in WT mice resolves but does not in Thy-1 -/- mice. When crossed with IL-1RI -/- mice, the double knockout Thy-1/IL-1RI mice exhibited diminished fibrosis. Altogether, these data demonstrated that IL-1β promotes the loss of Thy-1 and the IL-1R signaling axis for the immuno-stromal crosstalk in pulmonary fibrosis.

## Methods

### Human Tissue Procurement

Human deidentified IPF lung samples were acquired from lung tissue explants of patients undergoing lung transplantation at the University of Minnesota. Excised tissue samples were flash frozen in OCT and stored at -80°C. Serial cryosections were acquired for histology and immunofluorescence.

### Mice and Intratracheal Bleomycin Administration

WT, Thy-1^-/-^, IL-1RI^-/-^, Thy-1^-/-^ IL-1RI^-/-^, and *Rosa26-CAG-ASC-mCitrine* mice on a C57BL/6 background were used between the ages of 10-14 weeks old for the bleomycin model. IL-1RI^-/-^ and ASC-Citrine mice were obtained from Dr. Sarah Ewald. Mice were anesthetized with an intraperitoneal ketamine/xylazine cocktail injection (60-80/5-10 mg/kg), suspended vertically on an angled stand, an angiocatheter inserted into the trachea, and 50μL of bleomycin (1U/kg) dissolved in sterile saline was administered with 50μL of saline as control. Mice were maintained in the UVA animal facility in accordance with guidelines established by IACUC and all experiments were in accordance with protocols approved by UVA IACUC.

### Tissue Harvesting and Cell Dissociation

For single cell dissociation of lungs after bleomycin administration, mice were euthanized at day 7 via carbon dioxide asphyxiation. The thoracic cavity was opened, the right ventricle of the heart was perfused with PBS to clear blood, and the lungs were excised. Lungs were finely minced with scalpels, moved to a digestion media (PBS + 0.05% Trypsin + 0.04% DNase + 0.1% Collagenase + 25μM HEPES) for 30 minutes at 37°C with mixing via pipetting every 15 minutes, and carefully strained through a 100 μm cell strainer. Cells were washed twice with PBS and ready for cell surface staining. For histology, mice were euthanized at days 14, 28, and 42 via carbon dioxide asphyxiation. The thoracic cavity was opened, the right ventricle of the heart was perfused with PBS to clear blood, lungs were filled intratracheally with OCT, and the lungs were excised and then submerged in OCT over dry ice. Cryoblocks were stored at - 80°C.

### Immunofluorescence Imaging and Histology

10um sections of IPF biopsy cores and mouse lungs were cut using a CryoStar NX50 cryostat and mounted on SuperFrost Plus slides for downstream histology and immunofluorescence. For immunofluorescence, cryosections were fixed with 4% paraformaldehyde and blocked with 3% normal goat serum (NGS) + 0.1% Triton in 1XPBS for intracellular antigens. Primary antibody dilutions were prepared in 3% NGS in 1XPBS as follows: anti-mouse α-SMA (1A4, ThermoFisher) at 1:200 and biotinylated anti-mouse IL-1RI (JAMA-147, BioLegend) at 1:100. The primary antibody incubation was done overnight at 4°C, then three 1XPBS washes, and then a 1-hour secondary antibody staining at room temperature. The secondary antibodies (goat anti-mouse Alexa Fluor 488 and Streptavidin conjugated with Alexa Fluor 555; Invitrogen) were all prepared in 3% NGS in 1XPBS at a dilution of 1:1000. DAPI was used as a counterstain at 300nM in 1xPBS for 5 minutes at room temperature and then mounted in mounting medium ProLong Diamond Antifade (ThermoFisher). Slides were imaged on a Keyence BZ-X810 microscope using the 10X objective.

### Conventional and Imaging Flow Cytometry

Fibroblasts were lifted off of hydrogels using TrypLE Express Enzyme (Thermo Fisher), washed with FACS Buffer (PBS + 5% FBS + 0.1% sodium azide), stained with the Zombie Near-Infrared (NIR) Fixable Viability dye (BioLegend) at 1:500, and then fixed using the FIX & PERM Cell Permeabilization Kit (ThermoFisher). Cell surface staining was done using the following antibodies: APC-conjugated anti-Thy-1.2 (clone: 53-2.1, BioLegend) at 1:100, purified anti-activated αvβ3 antibody (clone: WOW-1) at 1:50, biotinylated anti-mouse IL-1RI (clone: JAMA-147, BioLegend) at 1:100, and TruStain FcBlock anti-mouse CD16/32 (clone: 93, BioLegend) at 1:50. The secondary antibodies (goat anti-mouse Alexa Fluor 488 and Streptavidin conjugated with Alexa Fluor 555; Invitrogen) were all prepared in FACS Buffer at a dilution of 1:1000. After surface staining, intracellular staining was done using the permeabilization buffer in the FIX & PERM kit with PE-conjugated anti-mouse α-SMA (clone: 1A4, R&D Systems) at 1:20 in the permeabilization buffer. Cells were then washed and analyzed on a BD LSR Fortessa through the UVA Flow Cytometry Core Facility and analysis done using FCS Express 7. For image flow cytometry, single cell suspension of bleomycin treated lungs were collected as described above. Cells were washed with FACS Buffer, stained with the Zombie NIR Viability dye at 1:500, and then fixed using the FIX & PERM kit. Cell surface staining was done with the following antibodies: BV421-conjugated anti-mouse CD45 (clone: 30-F11, BioLegend), BV605-conjugated anti-mouse EpCAM/CD326 (clone: G8.8, BD Biosciences), PE/Dazzle 594-conjugated anti-mouse CD31 (clone: 390, BioLegend), PE-conjugated anti-mouse Ly6G (clone: 1A8, BioLegend), BV605-conjugated anti-mouse CD3 (clone: 17A2, BioLegend), BV480-conjugated anti-mouse CD11b (clone: M1/70, BD Biosciences), APC-conjugated anti-mouse CD19 (clone: 1D3/CD19, BioLegend), PE/Dazzle 594-conjugated anti-mouse CD11b (clone: M1/70, BioLegend), and APC-conjugated anti-mouse CD11c (clone: N418, BioLegend). Cells were then washed with FACS Buffer and analyzed on a Amnis ImageStreamX Mark II image flow cytometer through the UVA Flow Cytometry Core Facility and analysis done using IDEAS software.

### Cell Culture

Mouse lung fibroblasts (MLFs) were cultured from digested mouse lungs in DMEM + 10% fetal bovine serum + 1% penicillin/streptomycin on tissue culture plastic for 1-2 weeks. Afterwards, MLFs were moved to 4kPa polyacrylamide hydrogels coated with gelatin (20μg/mL) to reduce the myofibroblastic phenotype that results from cells cultured on tissue culture plastic. After 1 week on gelatin-coated hydrogels, MLFs are lifted using Trypsin-EDTA (0.25%) and transferred to 4kPa hydrogels coated with fibronectin (10μg/mL) for experiments.

### Statistics

Statistics were performed using GraphPad Prism 9 (GraphPad, San Diego, CA). For multiple group comparisons, we performed 1-way ANOVA followed by Tukey’s post hoc test. When comparing two groups, a Student’s t-test was used. The number of mice used for each analysis was determine by power analysis (power = 0.80, α = 0.05). Statistical significance was defined at p-values < 0.05. All data are presented as mean ± SEM

## Results

Given the elevated levels of IL-1β reported in the fibrotic lungs of mice and IPF patients [14], we began by asking whether IL-1R expression was present within the fibrotic lung and where it was localized. Using tissue sections of lung biopsies from IPF patients, we observed IL-1R expression within the fibroproliferative regions of the IPF lung and those same regions exhibited robust αSMA expression (Fig 1). These data indicate that inflammatory signaling is present in the regions of the fibrotic lung undergoing active fibrosis, and that IL-1R expression is localized to these regions.

**Figure 1.**
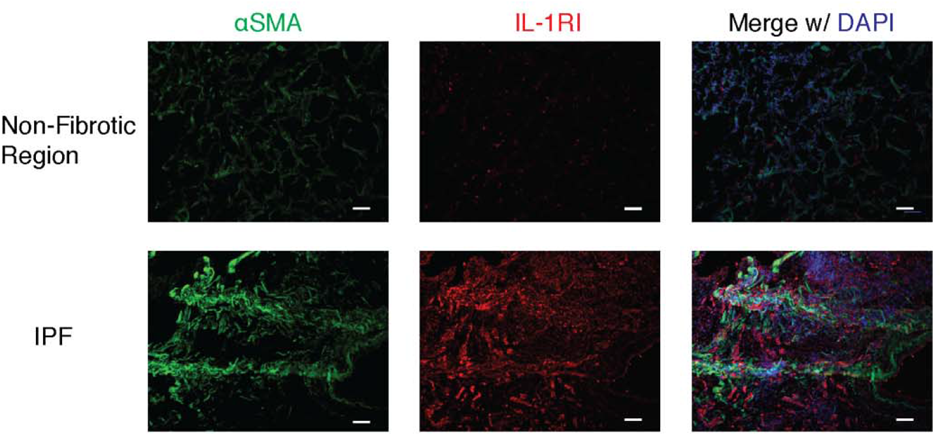
IL-1RI expression in IPF lung tissue. Representative images of immunofluorescence of αSMA and IL-1RI. OCT-embedded IPF lung biopsies were cryosectioned at 10μm, mounted, stained for αSMA to identify fibroblastic foci, and stained for IL-1RI as well to determine expression among fibroblastic foci. Sections were counterstained with DAPI. Scale bars are 100 μm.

In order to determine the functional relevance of IL-1R to lung fibrosis, we used the bleomycin-induced lung fibrosis model in WT and IL-1R -/- mice. Based on H&E and Picrosirius Red staining, bleomycin-induced fibrosis was dramatically diminished among IL-1R-deficient mice, further implicating IL-1R signaling in lung fibrosis (Fig 2). Identifying the cell population releasing the IL-1R ligand IL-1β during lung fibrosis is challenging for several reasons. First, IL-1β is stored as an inactive a precursor (pro-IL-1β) and is cleaved into its mature form by caspase-1 via inflammasome activation so transcriptional upregulation is not a proxy for functional IL-1 release [22]. Second, IL-1β is released by inflammasome-mediated pore formation and rapid inflammatory cell death making these cells difficult to identify *in situ* [23]. To circumvent challenges and identify which cell populations predicted to release IL-1β we applied the bleomycin-induced fibrosis model in mice expressing a transgene for apoptosis-associated speck-like protein containing a caspase recruitment domain (ASC) tagged with the fluorescent protein mCitrine (Figure 3A). During inflammasome activation, ASC oligomerizes with inflammasome sensors and caspase-1, forming a macromolecular complex that can be visualized by microscopy that catalyzes IL-1β maturation and release (Figure 3B) [22,24]. On day 7 after mice received intratracheal bleomycin delivery to induce pulmonary fibrosis, lungs were harvested and dissociated for analysis on the ImageStream imaging flow cytometer to identify the cell population(s) that formed ASC puncta in their cytosol. Using the “Spot Count” feature in the IDEAS software ASC speck formation on immune cells (CD45), endothelial cells (CD31), and epithelial cells (EpCAM/CD326) and cells lacking all three markers, considered the stromal compartment, was assessed (Figure 3C). ASC speck formation was most abundant on the CD45+ CD11b myeloid cells and to a lesser extent Ly6G+ neutrophils and CD3+ T cells indicating that several immune compartment cell types may be able to initiate IL-1R signaling in the lung during bleomycin-induced fibrosis (Fig 3D).

**Figure 2.**
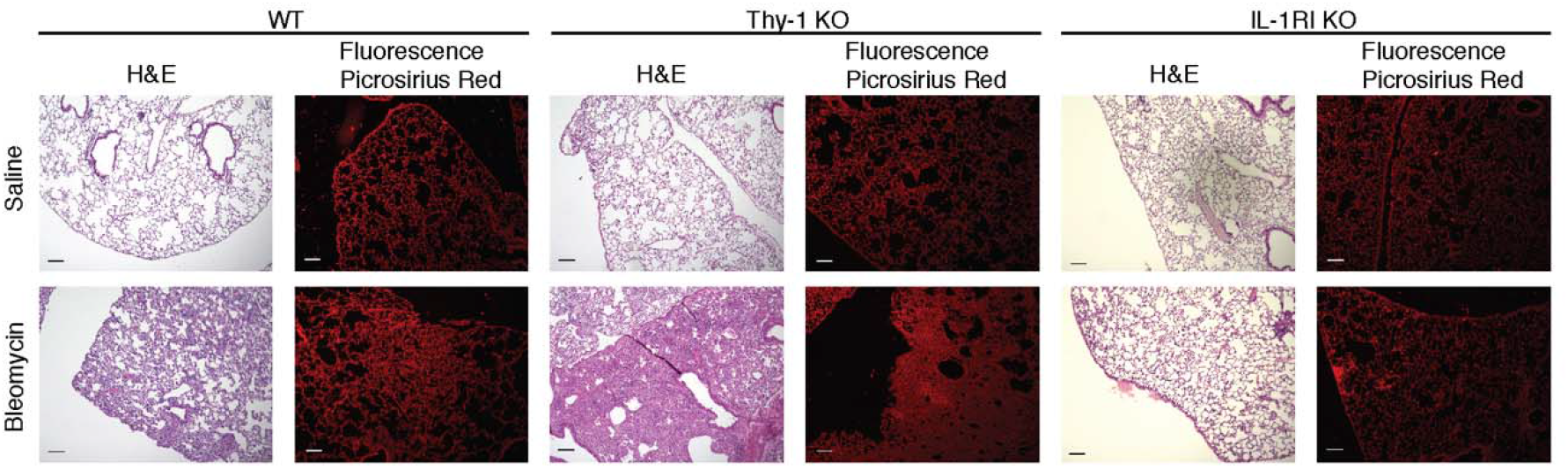
Loss of IL-1R signaling ameliorates bleomycin-induced lung fibrosis. Histological analysis of bleomycin-induced pulmonary fibrosis in WT, Thy-1 KO, and IL-1RI KO mice using H&E and picrosirius red staining among mice sacrificed on day 28. Changes in collagen content were quantified using picrosirius red micrographs and images were analyzed using Fiji. Scale bars are 100 μm. n=5 mice with 5 images per slide. *p<0.05.

**Figure 3.**
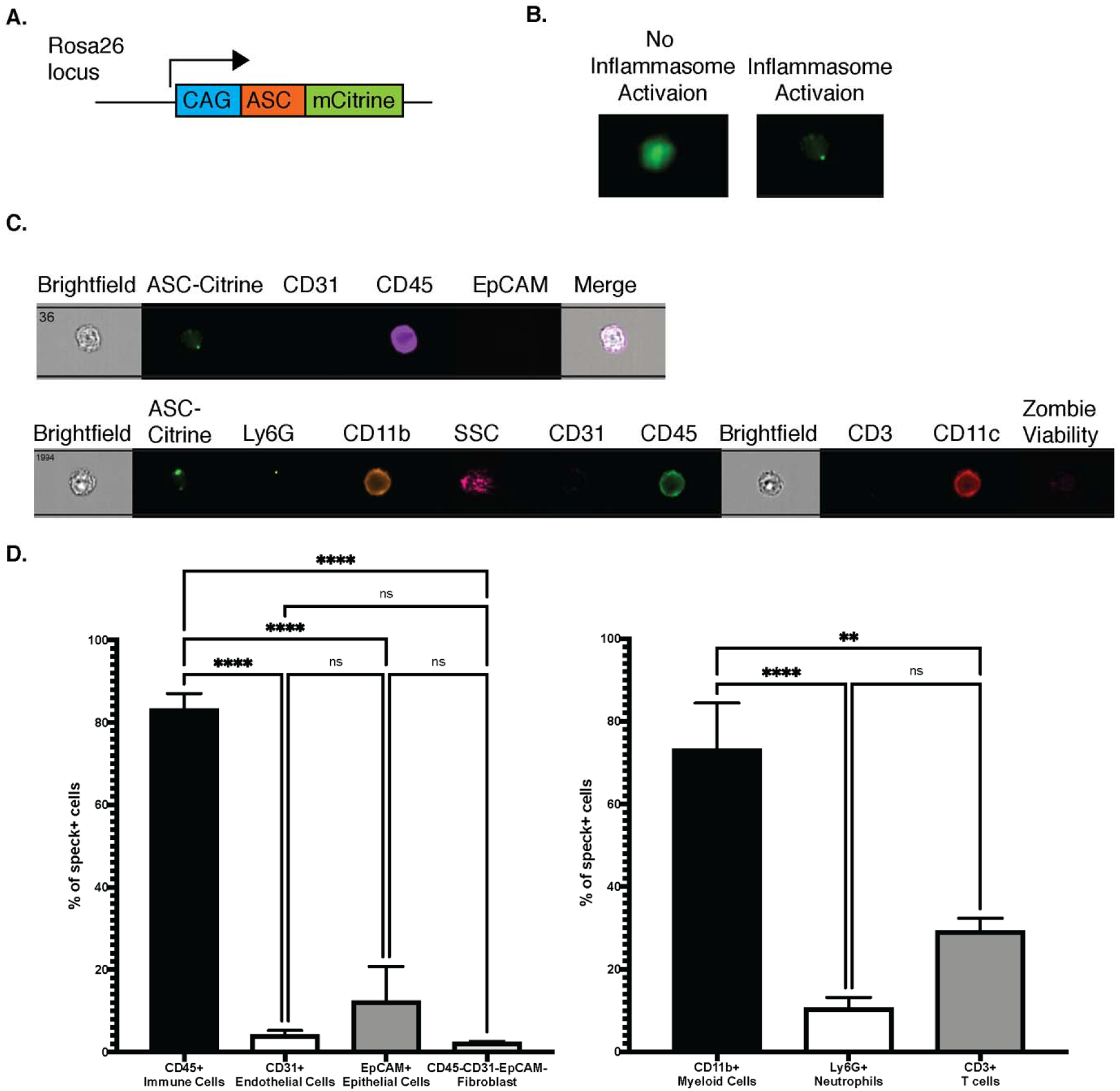
Myeloid cell inflammasome activation via ASC speck formation during lung fibrosis. A) Fluorescent reporter mice with mCitrine fused to the inflammasome related protein ASC on the *Rosa26* locus were treated with bleomycin and sacrificed on day 7 for imaging flow cytometry. B) With no inflammasome activation, Citrine distribution was evenly throughout the cell. However, during inflammasome activation ASC helps to create the macromolecular inflammasome complex, which appears as a speck with mCitrine fused to ASC. C) Using the ASC-mCitrine mice in the bleomycin model, immune cells were the dominant cell population demonstrating ASC speck formation. Among immune cells, CD11b+ myeloid cells were the dominant cell population demonstrating ASC speck formation based on D) the quantification and characterization of speck positive cells. n=3-5 mice where the mean of a mouse is averaged with all mice per group. *p<0.05, **p<0.01, ***p<0.001, ****p<0.0001

IL-1R signaling has long been implicated in lung fibrosis, yet what has remained unclear is the role of this inflammatory pathway in the pathogenesis of the disease. Previous work from our group and others have demonstrated the loss of Thy-1, a GPI-anchored membrane protein, in pulmonary fibrosis. Thy-1 facilitates normal mechanotransduction by regulating αvβ3 integrin conformation in response to substrate stiffness [10]. αvβ3 integrin activation has been shown to promote cell contractility and strain stiffening of the local microenvironment [7]. However, it is unknown what promotes the loss of Thy-1. We hypothesized that elevated proinflammatory cytokines such as IL-1β promote the loss of Thy-1 and lead to the emergence of Thy-1 negative fibroblasts observed in the bleomycin model [25] and in IPF [10]. We tested this hypothesis by seeding mouse lung fibroblasts on fibronectin-coated 2kPa polyacrylamide hydrogels and treating fibroblasts with IL-1β and measure both Thy-1 expression and αvβ3 integrin activation. In response to IL-1β treatment, mouse lung fibroblasts downregulated surface Thy-1 expression and upregulated expression of αvβ3 integrin in the active conformation stained by the conformation sensitive WOW-1 antibody (Fig 4A). As expected, when gating on Thy-1, active αvβ3 was expressed at higher levels on Thy-1 negative compartment while on soft 2kPa hydrogels. Unexpectedly, expression of the IL-1β receptor, IL-1RI, was also upregulated on the Thy-1 negative compartment (Fig 4B). IL-1β is considered an alarmin, and the stringent regulation of IL-1β expression, maturation, and release is thought to play an important role in limiting the magnitude of the IL-1 response in local tissues, it follows that cell type specific upregulation of IL-1R1 could dramatically increase the sensitivity of the Thy-1 negative fibroblast population in the tissue microenvironment. This indicates that Thy-1 negative fibroblasts are a unique subpopulation poised to respond to inflammatory cytokines, such as IL-1β, and also contribute to changes in mechanotransduction.

**Figure 4.**
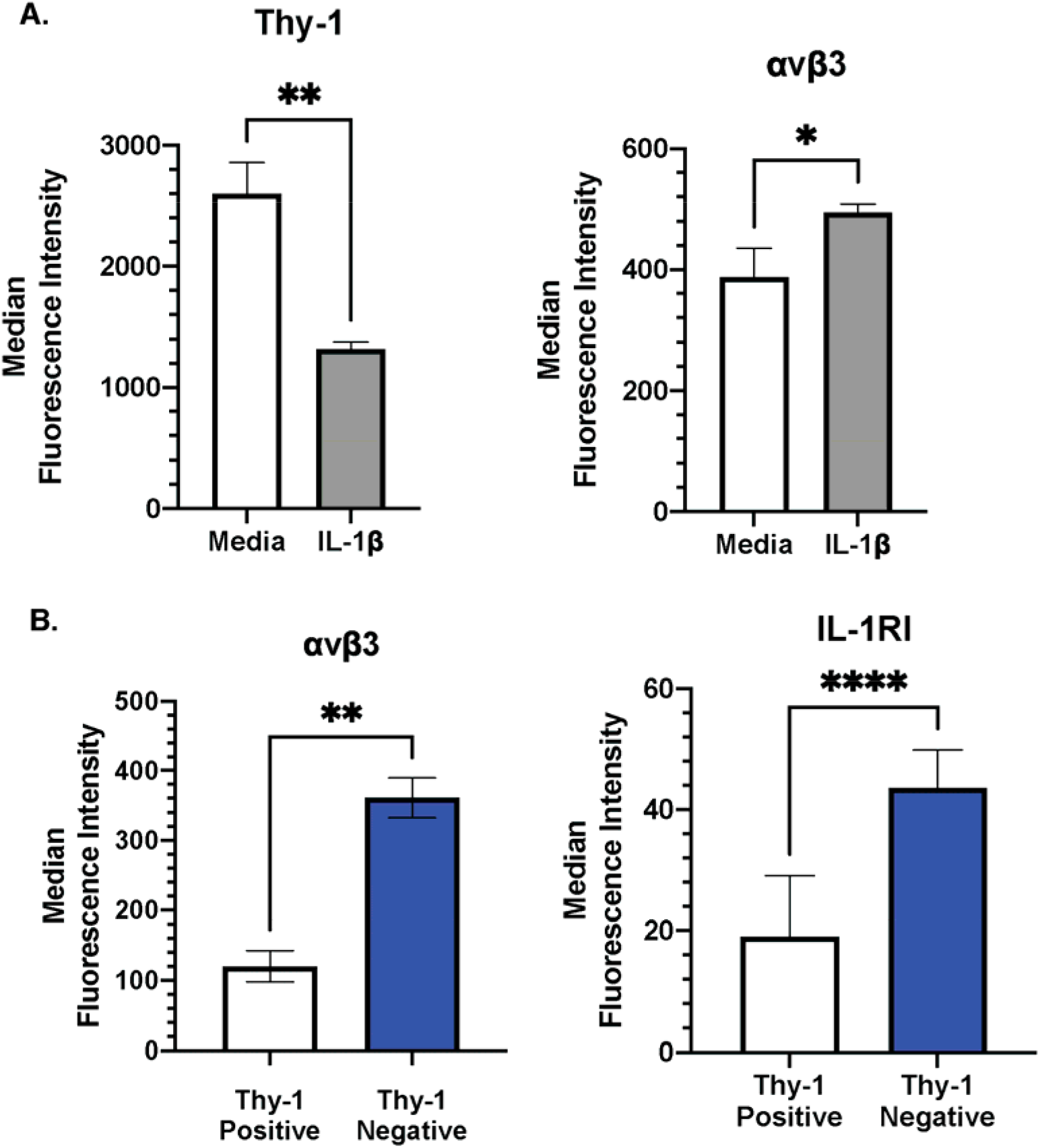
IL-1β treatment among MLFs on soft hydrogels prompted the emergence of an inflammatory subpopulation with dysfunctional mechanotransduction. A) Mouse lung fibroblasts (MLFs) were treated with IL-1β for 72 hours while on soft fibronectin-coated 2 kPa polyacrylamide hydrogels. IL-1β treatment promoted Thy-1 loss measured via flow cytometry. Additionally, while on soft hydrogels, IL-1β treatment among MLFs elevated αvβ3 integrin activation measured via flow cytometry using the WOW-1 antibody. B) Among Thy-1 negative MLFs, αvβ3 integrin activation was elevated. Surprisingly, IL-1RI expression was elevated as well among Thy-1 negative MLFs. (n=5). *p<0.05, **p<0.01, ***p<0.001, ****p<0.0001.

To determine whether the immune-responsive quality of Thy-1 negative fibroblasts contributed directly contributed to pulmonary fibrosis, we leveraged previous work in our group that demonstrated that Thy-1^-/-^ mice, unlike WT mice in the bleomycin pulmonary fibrosis model, demonstrated non-resolving fibrosis. Typically, bleomycin-induced pulmonary fibrosis peaks by day 14 and resolves by day 42 in WT mice. However, in Thy-1^-/-^ mice, fibrosis continues to worsen out to day 42[7] (Figure 5 left and center). To determine whether non-resolving fibrosis driven by the Thy-1 deficiency was dependent on the IL-1R signaling axis, we generated Thy-1^-/-^ IL-1RI^-/-^ mice. The absence of IL-1R limited the severity of acute bleomycin induced fibrosis the non-resolving nature of disease (Fig 5). This indicates that the IL-1 signaling axis is necessary for the robust fibrotic phenotype observed in the absence of Thy-1.

**Figure 5.**
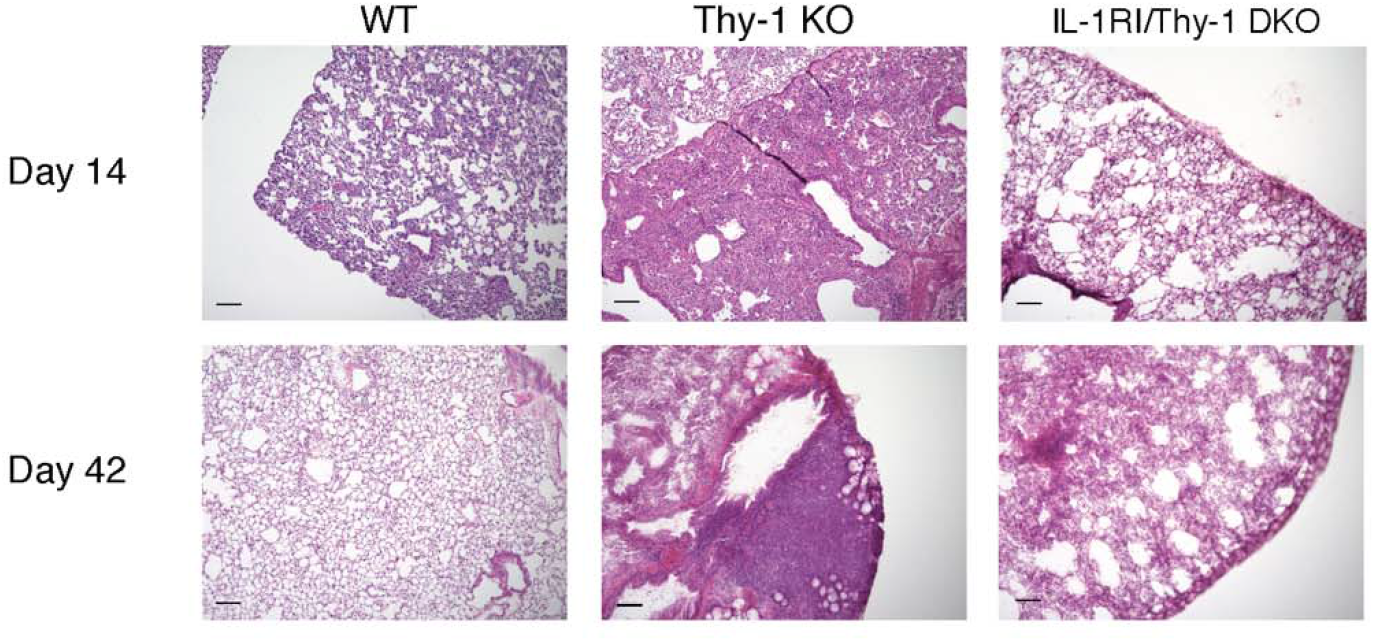
Loss of IL-1R signaling on a Thy-1 deficient background reduced lung fibrosis. WT mice demonstrate fibrosis resolution by day 42, whereas Thy-1 KO mice exhibit non-resolving fibrosis in response to intratracheal bleomycin. When IL-1RI/Thy-1 double KO mice receive intratracheal bleomycin, the loss of IL-1 signaling reduces non-resolving fibrosis. Additionally, Thy-1 negative fibroblasts lose elevated IL-1RI, whose loss corresponds to reduced lung fibrosis. (n=5 mice per group).

## Discussion

The current understanding of fibrosis progression is characterized by dysfunctional fibroblast mechanotransduction (i.e. sensing a soft tissue as stiff), increased tissue stiffness mediated by myofibroblasts laying down more ECM, and myofibroblasts contracting the surrounding tissue [7]. This creates a positive feedback loop that stiffens more surrounding tissue and promotes the recruitment of naïve fibroblast into this loop thus driving the pathology of IPF. However, what we have not understood for a long time is how all of this is initiated. How is soft tissue perceived as stiff? This dysfunctional mechanotransduction indicates the presence of a pro-fibrotic lung fibroblast population in the foci that emerges prior to the stiffening of lung tissue, indicating an early stiffness-independent pathway. If fibrosis is initiated on soft ECM, it is important to understand the mechanism by which that happens within the foci. Previous work demonstrated that the fibroblastic foci is characterized by Thy-1 (-), αSMA+ myofibroblasts that are within foci [6]. Further work determined that Thy-1 contributes to rigidity sensing and mechanotransduction through its binding to inactive αvβ3 integrin on soft (∼2kPa) substrates, a stiffness similar to healthy lung tissue, and enabling active αvβ3 integrin on stiff (∼20kPa) substrates, a stiffness similar to fibrotic lung tissue [10]. A question that had not been answered yet is what promotes the loss of Thy-1. Elevated inflammation and aberrant mechanotransduction have been observed in various fibrotic disorders, yet there has remained a critical gap between the two that is necessary to bridge in order to better understand the pathogenesis of pulmonary fibrosis, but also other fibrotic disorders as well. Here we demonstrate that IL-1β treatment promotes the loss of Thy-1, and this loss corresponds to aberrant αvβ3 integrin activation while on soft hydrogels. Furthermore, this loss confers on those fibroblasts an immuno-responsive phenotype based on elevated IL-1RI expression. This fibroblast subpopulation offers a promising theory as to how inflammation bridges to tissue remodeling and fibrosis, beyond simply soluble factors being sent from immune cells to stromal cells. But rather that specific subpopulations of cells mediate this transition from inflammation to tissue remodeling and that the emergence of this immuno-responsive fibroblast subset seems to contribute to pulmonary fibrosis. We showed that in the αSMA rich regions of IPF tissue, αSMA+ myofibroblasts co-express IL-1RI, implicating them in human disease. Furthermore, when Thy-1 (-) fibroblasts lacked IL-1 signaling via the Thy-1^-/-^IL-1RI^-/-^ mouse, non-resolving fibrosis observed in Thy-1^-/-^ mice was diminished. This subpopulation and the IL-1 signaling axis offers a new potential explanation of the pathogenesis of pulmonary fibrosis, and the notion of an immuno-responsive stromal population might help us better understand other fibrotic diseases as well. Furthermore, targeting IL-1 signaling in foci offers a new therapeutic target that might help stop fibrosis progression in disease.

